# Efficient characterization of multiple binding sites of small molecule imaging ligands on amyloid-beta, 4-repeat/full-length tau and alpha-synuclein

**DOI:** 10.1101/2023.03.12.531651

**Authors:** Jens Sobek, Junhao Li, Benjamin F. Combes, Juan A Gerez, Peter K. Nilsson, Martin T. Henrich, Fanni F. Geibl, Kuangyu Shi, Axel Rominger, Wolfgang H. Oertel, Roger M. Nitsch, Agneta Nordberg, Hans Ågren, Roland Riek, Ruiqing Ni

## Abstract

**Aim:** There is an unmet need for compounds that detect alpha-synuclein (αSyn) and 4-repeat tau, which are critical in many neurodegenerative diseases for diagnostic and therapeutic purposes. Here, we aim to develop an efficient surface plasmon resonance (SPR)-based method to facilitate the characterization of small molecule ligands/compounds to these fibrils.

**Methods:** SPR measurements were conducted to characterize the binding properties of fluorescent ligands/compounds towards recombinant Aβ_42_, K18 4-repeat/full-length tau and αSyn fibrils. In silico modelling was performed to examine the binding pockets of ligands on αSyn fibrils. Immunofluorescence staining with fluorescence ligands and specific antibodies on postmortem brain tissue slices from patients with Parkinson’s disease and disease mouse models was performed.

**Results:** We optimized the protocol for immobilizing Aβ_42_, K18 tau, full-length tau and αSyn fibrils in a controlled aggregation state on SPR sensor chips. The results from the analysis of binding kinetics suggested the presence of at least two binding sites for all fibrils, including luminescent conjugated oligothiophenes (HS-169, HS-84, h-FTAA and q-FTAA), pyridine derivative PBB5, nonfluorescent methylene blue and lansoprazole. In silico modelling studies for αSyn (6H6B) showed four binding sites with preference to S4. Immunofluorescence staining validated the detection of pS129-positive αSyn in brain tissue from Parkinson’s disease patients, αSyn PFF-injected mice, 6E10-positive Aβ in arcAβ mice, and AT-8/AT-100-positive in tau pR5 tau mice, respectively.

**Conclusions:** SPR measurements of ligands and small molecules binding to Aβ_42_, 4R and full-length tau and αSyn fibrils suggest the existence of multiple binding sites. This approach may provide efficient characterization of compound binding properties towards these fibrils important in neurodegenerative diseases.

## 1 Introduction

Neurodegenerative diseases represent a tremendous unmet clinical need. A common feature of these diseases is the abnormal cerebral accumulation and spreading of pathological protein aggregates, affecting selective vulnerable circuits in a disease-specific pattern (Jucker and Walker, 2018; Riek and Eisenberg, 2016). Alzheimer’s disease (AD) is pathologically hallmarked by amyloid-β (Aβ) plaques and neurofibrillary tangles of hyperphosphorylated tau. Other tauopathies include frontotemporal dementia with 4R and 3R tau, progressive supranuclear palsy (PSP) with 4R tau, and corticobasal degeneration (CBD) with 3R tau accumulation (Spillantini and Goedert, 2013). α-synucleinopathy is characterized by the accumulation of alpha-synuclein (αSyn) in Parkinson’s disease (PD), dementia with Lewy bodies and multiple system atrophy (MSA). The use of positron emission tomography (PET) with Aβ and tau imaging ligands has facilitated the early/differential diagnosis of AD (Dubois et al., 2021). Currently, there is an unmet clinical need for PET ligands for 4-repeat tau and αSyn aggregates to assist in diagnostic and clinical outcome evaluations. Several imaging ligands are currently in the pipeline, such as [^18^F]ACI-12589 (Smith et al., 2022), [^11^C]MODAG-001 (Kuebler et al., 2021), [^18^F]SPAL-T-06 (Matsuoka et al., 2022) and [^18^F]UCB-2897 (NCT05274568). In addition, optical imaging ligands have been developed and applied in mechanistic and treatment studies using disease animal models recapitulating amyloidosis/tauopathy/αSyn. Using two-photon and diffuse optical imaging approaches, several imaging ligands, such as BTA-1, methoxy-X04, BF-158, PBB5, luminescent conjugated oligothiophenes (LCOs), BODIPY derivatives and fluorescently labelled antibodies, have been reported (Bian et al., 2021; Calvo-Rodriguez et al., 2019a; Hou et al., 2023; Krishnaswamy et al., 2014; Kuchibhotla et al., 2014; McMurray et al., 2021; Ni et al., 2020; Reyes et al., 2021; Tanriöver et al., 2020; Torre-Muruzabal et al., 2023; Vagenknecht et al., 2022; Verwilst et al., 2017; Wu et al., 2018). A number of experimental techniques, including fluorescence assays, radioligand competition assays (LeVine, 2005; Malarte et al., 2020; Ni et al., 2017; Ni et al., 2013a; Ni et al., 2018; Ni et al., 2013b; Ni et al., 2021a; Yap et al., 2021), nuclear magnetic resonance spectroscopy (Gerez et al., 2020; Schütz et al., 2018), and cryogenic electron microscopy (cryo-EM) (Antonschmidt et al., 2022; Shi et al., 2021a), have been reported to investigate interactions between ligands and Aβ peptides (Ferrie et al., 2020; Sanna et al., 2021; Shi et al., 2021b; Shimogawa and Petersson, 2021; Yang et al., 2022) and have demonstrated multiple ligand binding sites. *In silico* studies have also often been applied to study the interactions between ligands and proteinopathies/fibrils (Kuang et al., 2020; Kuang et al., 2019; Murugan et al., 2016; Murugan et al., 2018; Zhou et al., 2021).

Surface Plasmon Resonance (SPR) is the method of choice to study the kinetics of interactions for a wide range of molecular systems and has been widely used in pharmaceutical, biosensing, and biomolecular research (Erbaş and Inci, 2022; Karlsson, 2004). The SPR approach has been more commonly used in detecting Aβ and tau monomers in biological samples, such as blood or cerebrospinal fluid, as diagnostic biomarkers (Kim et al., 2016; Nangare and Patil, 2021; Rezabakhsh et al., 2020; Stravalaci et al., 2012). It has also been used to investigate Aβ elongation, aggregation dynamics (Ryu et al., 2008; Stravalaci et al., 2011) and interactions with aggregation inhibitors (De Simone et al., 2013; Dehghani et al., 2021; Frenzel et al., 2014; Marasco et al., 2021). A few studies have reported SPR for αSyn fibrils (Honarmand et al., 2019; Jha et al., 2017; Sangwan et al., 2020; Thom et al., 2021; Yin et al., 2020), tau fibrils (Haghaei et al., 2020; Lisi et al., 2017; Rojo et al., 2010), and Aβ fibrils (Martins et al., 2015; Rojo et al., 2010) for characterization of interactions with resveratrol and lansoprazole. However, binding kinetics and binding site determination need further optimization of the experimental protocol and modelling. Here, we established a SPR pipeline to determine the binding kinetics of small molecules (imaging ligands and nonfluoresence compounds) on Aβ_42_, K18 tau, full-length tau and αSyn fibrils. We further examined the binding sites of fluorescence ligands on αSyn fibrils by *in silico* modelling and immunofluorescence staining in postmortem human brain tissues from patients with PD and from PFF-injected mouse models.

## 2 Methods

### 2.1 Chemicals and antibodies

Luminescent conjugated oligothiophenes (LCOs), including HS-84, HS-169, h-FTAA, and q-FTAA, were synthesized and provided by KPRN (Linköping, Sweden) (Klingstedt et al., 2011; Shirani et al., 2015). Methylene blue (Sigma Aldrich, Switzerland), lansoprazole (Sigma Aldrich, Switzerland), and pyridine-derivative PBB5 (RadiantDye, Germany) were purchased from the indicated sources (chemical structures in **SFig. 1**). Detailed information on the chemicals and antibodies used in the study is provided in **STable 1**.

### 2.2 Recombinant Aβ_42_, K18 tau, full-length tau and αSyn fibril production, characterization and detection by fluorescence ligands

Recombinant Aβ_42_, K18 tau (4-repeat), full-length tau and αSyn were expressed and produced by *E. coli* as described previously (detailed methods in supplementary material) (Burmann et al., 2019; Gerez et al., 2019). As a substrate for HRP, Pierce ECL and Western Blotting Substrate (Thermo Fisher Scientific, U.S.A.) were used to detect Aβ_42_ and αSyn. To detect K18 tau, ECL Prime Western Blotting Detection reagents (Cytiva, U.S. A.) were used. Images of the resulting blots were taken with an ImageQuant LAS 4000 (GE Healthcare). After incubation of the protein solutions, fibrillization was verified by thioflavin T fluorescence assay: 45 μL of thioflavin T (5 μ either 2 μL of the incubated αSyn, 5 μ of K18 tau, or 5 μL of Aβ_42_ in a 45 µL quartz cuvette (quartz SUPRASIL Ultra Micro Cell, Hellma). Mass photometry was used to measure the mass of the fibrils at FGCZ, ETH Zurich.

Transmission electron microscopy (TEM) was performed by adding 4 μL of the fibril samples (∼50 μM) in PBS directly to the negatively glow-discharged carbon-coated copper grids, followed by incubation for 1 minute at room temperature. The excess solution was gently removed using Whatman filter paper. Samples were stained with 10 μL of an aqueous phosphotungstic acid solution (1%, pH 7.2) for 1 minute. The excess stain on the grid was then wiped off with filter paper, washed with double-distilled water and air dried. Finally, the images were recorded on a Morgagni 268 electron microscope (FEI GmbH, Germany) at ScopeM, ETH Zurich.

For fluorescence assays with different fluorescence-emitting ligands, mixtures of different fibrils and ligands were incubated for 1 minute and resuspended in a pipette, and the fluorescence intensity over a wide range of spectra was measured with a spectrofluorometer (FluoroMax-4, Horiba Jobin Yvon) using a known excitation wavelength for these ligands.

### 2.3 Surface Plasmon Resonance

SPR measurements were conducted in HBS buffer at 20 °C using Biacore instruments T200 and S200 (Cytiva, Uppsala, Sweden). Aβ_42_, tau, and αSyn recombinant fibrils were immobilized in the flow cell of sensor chips produced by Xantec (Düsseldorf, Germany) and Cytiva by coupling fibril amine groups to sulfo-N-hydroxysuccinimide (sulfo-NHS)-activated carboxylic acid groups on the chip surface. Since no nonbinding fibril(s) were available, the reference flow cell remained empty. Different chip surfaces were tested to obtain a high immobilization density and low nonspecific adsorption of analytes, including carboxymethylated dextran surface CM5 (Cytiva, Sweden), carboxymethyldextran hydrogel surface CMD200M, linear polycarboxylate hydrogel surface HC30M and HC1500M, linear polycarboxylate hydrogel (reduced charge) surface HLC30M, and zwitterionic hydrogel surface ZC150D (all from Xantec, Germany). All steps for surface preparation and immobilization were conducted at a flow rate of 5 µL/min. Surface carboxylic acid groups were activated with 0.2 M sulfo-NHS and 1-ethyl-3-(3-dimethylaminopropyl)carbodiimide (EDC) (both from Xantec) 0.1 M (CM5 and CMD200M: 0.05 M) in 10 mM 2-(N-morpholino)ethanesulfonic acid (MES buffer, abcr Swiss AG, Switzerland) at pH 5.0 for 550 s (CM5 and CMD200M: 180 s) using 5 mM MES pH 6.5 as running buffer. K18 tau and αSyn fibrils were diluted to 2.5-10 µM in 5 mM acetate buffer (Fisher Scientific, Switzerland) at pH 4.5 and coupled for 750 s at a flow rate of 5 µL/min on all surfaces except for ZD150D. Surfaces were blocked with 1 M ethanolamine (Xantec, Germany) for 500 s. On the ZC150D chip, K18 tau, αSyn and Aβ_42_ were immobilized at 2.5 – 10 µM in 5 mM MES pH 6.5 for 750 s to a surface activated with 500 mM EDC and 100 mM sulfo-NHS for 600 s in 5 mM MES pH 6.5. Finally, the surface was blocked with 1 M glycine in 10 mM MES pH 6.5.

Binding kinetics were measured after a stabilization period of a few hours that was necessary due to a slightly decreasing baseline presumably caused by a slow dissociation of fibrils. Immobilized fibrils were stable for a few days. Kinetic constants for binding were determined by injection of a dilution series of 5-8 concentrations of analyte in HBS buffer (Teknova, USA) at a flow rate of 30 µL/min in duplicate (“full kinetics”). Since no regeneration conditions were found that did not lead to decomposition of the fibrils, single cycle kinetics (Karlsson et al., 2006) were measured in cases of slow dissociation by consecutive injection of 5 analyte concentrations followed by a dissociation period of 10 min.

Data were evaluated using BiaEvaluate v2.03 software (Cytiva, Sweden). Binding kinetics were analysed globally using different kinetic models, including 1+1, a sum of 2 exponentials model (“heterogeneous ligand” in terms of BiaEvaluate), and a sum of three exponentials model that was created using the script editor of the software. The software does not allow the creation of higher-order kinetic models.

### 2.4 In silico modelling and calculations

The αSyn fibrils were adopted from previous studies (PDB codes: 6H6B (Guerrero-Ferreira et al., 2018; Kumari et al., 2021) and 2N0A (Kuang et al., 2019)). The protein preparation wizard module in the Schrödinger suite (V. 2021-3, Schrödinger LLC, New York, U.S.A.) was used to add hydrogen atoms and determine the protonation states of the ionizable residues. The initial structures of the tested compounds were drawn in the Maestro interface and optimized using the Ligprep module (Schrödinger Release 2021-3: LigPrep, Schrödinger, LLC, New York, U.S.A.). The Glide module was used for docking, in which we used normal inner and outer box sizes of 10 and 20 Å, respectively (Friesner et al., 2004; Halgren et al., 2004). The standard precision (SP) mode was used for dockings, while other settings were left as default in Glide. The top 1 scored binding pose of each ligand–αSyn complex was subjected to molecular dynamics (MD) simulations. All MD simulations were carried out for 100-ns production with the OPLS-4 force field (Lu et al., 2021) using the Desmond module. In each simulation, the protein[ligand complex was centered into an orthorhombic box with a boundary buffer of 12 Å, and ∼24 000 TIP3P water (Jorgensen et al., 1983) molecules and counter ions were added to neutralize the system. Additional Na^+^ and Cl^-^ were added to reach a 0.15 M salt concentration. Before the production simulations, the system is energy minimized and equilibrated with a Nose [ Hoover chain thermostat (300.0 K) (Martyna et al., 1992) and Martyna-Bobias-Klein barostat (1.0 atm) (Martyna et al., 1994) using the default protocol implemented in Desmond. Molecular mechanics (MM)/generalized born solvent accessibility (GBSA)) (Kollman et al., 2000) binding free energy for each ligands was averaged from a total of 200 snapshots evenly extracted from the 100-ns trajectories. The Prime module [Schrödinger Release 2021-3: Prime, Schrödinger, LLC, New York, NY, 2021.] was used for the MM/GBSA calculations. The protein[ligand complexes were refined and optimized using the OPLS4 force field with the VSGB (variable dielectric surface generalized Born) continuum solvation model (Li et al., 2011). For the minimization, the residues within 5 Å of the ligand were included. After that, the MM/GBSA method implemented in the Prime module was used to rescore the binding poses. Root-mean-square deviation (RMSD) analysis of the average distance between a group of atoms (e.g., backbone atoms of the four binding sites on αSyn) was performed.

### 2.5 Postmortem human brain tissue

Three PD cases (mean age 72.3[9.5]) each with a clinical diagnosis confirmed by pathological examination of Lewy bodies (Braak LB 6, with tau Braak 0-1 and Aβ Ο), and one nondemented control aged 69 y (Braak LB 0, with tau Braak 1 and Aβ −), were included in this study (detailed information in **Table 1**). Paraffin-embedded autopsy brain tissues from the medulla oblongata and cerebellum with high αSyn inclusion accumulation were obtained from the Netherlands Brain Bank (NBB), Netherlands. All materials had been collected from donors or from whom written informed consent for a brain autopsy and the use of the materials and clinical information for research purposes had been obtained by the NBB. The study was conducted according to the principles of the Declaration of Helsinki and subsequent revisions. All experiments on autopsied human brain tissue were carried out in accordance with ethical permission obtained from the regional human ethics committee in Canton Zurich and the medical ethics committee of the VU Medical Center for the NBB tissue.

**Table 1.** Information on brain tissue samples from patients with Parkinson’s disease, nondemented control and animal models.

| Postmortem human brain sample |  |  |  |  |  |  |  |  |
| --- | --- | --- | --- | --- | --- | --- | --- | --- |
| No | Sex | Age (y) | PM delay (h) | Braak tau | Amyloid-β | Braak LB | Diagnosis | Region |
| 1 | M | 61 | 7.5 | 1 | O | 6 | PD | MO |
| 2 | M | 65 | 4.8 | 0 | O | 6 | PD | MO |
| 3 | M | 83 | 5.3 | 1 | O | 6 | PD | LC |
| 4 | F | 69 | 8.5 | 1 | - | 0 | NC | MO |
| Mouse brain sample |  |  |  |  |  |  |  |  |
| Model |  | Number | Sex | Age (m) | Region imaged |  |  |  |
| αSyn PFF-injected mice | | 2 | M | 5.5 | CeA, NAc, PPN, PAG, SNc | | | |
| arcAβ mice | | 2 | M/F | 18 | Ctx | | | |
| pR5 mice |  | 2 | M/F | 18 | Hip |  |  |  |
| Nontransgenic littermate mice * |  | 4 | M/F | 18 | Ctx, Hip |  |  |  |

Ctx, Cortex; CeA, central amygdala; LB, Lewy body; LC; MO, medulla oblongata; NAc, nucleus accumbens; NC, nondemented control; PD, Parkinson's disease; PM, postmortem; PPN, pedunculopontine nucleus; PAG, periaqueductal gray; SNc, substantia nigra pars compacta; \* Two nontransgenic littermate mice for arcA $\beta$ and pR5 mice, respectively.

### 2.6 Animal models

Two transgenic arcA β mice overexpressing the human APP695 transgene containing the Swedish (K670N/M671L) and Arctic (E693G) mutations under the control of the prion protein promoter and two age-matched nontransgenic littermates of both sexes (18 months of age) (Knobloch et al., 2007; Ni et al., 2021b; Ren et al., 2022). Two transgenic MAPT P301L mice overexpressing human 2N/4R tau under the neuron-specific Thy1.2 promoter (pR5 line, C57B6. Dg background) and two age-matched nontransgenic littermates of both sexes (18 months of age) (Gotz et al., 2001; Massalimova et al., 2021; Vagenknecht et al., 2022). For the αSyn PFF mouse model, two male C57BL/6J mice (Charles River, Sulzfeld, Germany), 8 weeks old at the beginning of the experiment, were used. Animals were housed in individually ventilated cages inside a temperature-controlled room under a 12-hour dark/light cycle with ad libitum access to food and water. For arcAβ and pR5 mice, all experiments were performed in accordance with the Swiss Federal Act on Animal Protection and approved by the Cantonal Veterinary Office Zurich (permit number: ZH162/20). For αSyn PFF mice, procedures performed were in accordance with the ethical standards of the institution at which the studies were conducted (Regierungspräsidium Giessen, GermanyV54–19c2015h01GI20/28). For induction of αSyn pathology in a total volume of 550 nl of αSyn, PFFs with a concentration of 2.5 µg/µl were stereotactically injected in the pedunculopontine nucleus (PPN) as described (Henrich et al., 2020). Twelve weeks postinjection, mice were sacrificed, and tissue was prepared for immunohistochemical analysis. arcAβ, pR5 and PFF injected mice were perfused under ketamine/xylazine (75/10/2 mg/kg or 50/4.5 mg/kg body weight*, i.p.* bolus injection) with ice-cold 0.1 M phosphate-buffered saline (PBS, pH 7.4) and 4% paraformaldehyde (PFA) in 0.1 M PBS (pH 7.4). After perfusion, the mice were decapitated, and the brains were quickly removed and fixed for 1 or 3 days in 4% PFA (pH 7.4) and 3 days in 30% sucrose solution. Brains were then frozen on dry ice and stored at -80 °C until sectioning.

### 2.7 *Ex vivo* immunofluorescence and confocal imaging

The details on the antibodies and ligand concentration are described in **STable 1**. Human paraffin-embedded brain sections (3 µm) were deparaffinized and rehydrated prior to an antigen retrieval step (citrate buffer pH 6.0) in a microwave for 20 min at 98 °C. The sections were then costained with anti-αSyn (phospho-S129, pS129) antibody plus fluorescent secondary antibody. Ligands were incubated after secondary antibody for 30 min. Sections were counterstained using 4’,6-diamidino-2-phenylindole (DAPI). Mouse brains were embedded in tissue freezing media (OCT Compound, Tissue Tek, USA) and cut into 30-40 μm thick consecutive coronal sections using a cryostat microtome (CM3050 S, Leica, Germany). All sections spanning the complete rostro-caudal extent of the brain were kept in the correct order and stored at 4 °C in cryoprotect-solution (1:1:3 volume ratio of ethylenglycol, glycerol and 0.1 M PBS) until further processing (Henrich et al., 2018). Anti-αSyn pS129 antibodies, anti-phospho-tau antibodies (AT-8, AT-100), or anti-Aβ_1-16_ antibody 6E10, plus fluorescent secondary antibody were used as previously described on brain slices from αSyn PFF-injected mice, arcAβ mice (Ni et al., 2022), and pR5 mice (Vagenknecht et al., 2022). Ligands were incubated after secondary antibody for 30 min. Sections were counterstained using DAPI. The brain sections were imaged at 20× magnification using an Axio Observer Z1 microscope (whole brain slide) and at 63× magnification using a Leica SP8 confocal microscope (Leica, Germany) for colocalization. Lambda scans were performed using a Leica SP8 confocal microscope for the emission spectrum of the ligands on the staining as previously described to validate the signal (Ni et al., 2020). The images were analysed using Qupath (Bankhead et al., 2017) and ImageJ (NIH, U.S.A).

## 3 Results

### 3.1 *In vitro* fluorescence binding assays in recombinant fibrils

We produced Aβ_42_, K18 tau, full-length tau and αSyn fibrils using bacterially produced recombinant monomers. During the processing of K18 tau, full-length tau and αSyn after each step, an aliquot was taken, and at the end, the identity was verified by SDS–PAGE (**SFig. 2**). The monomers were validated using western blotting, and the fibrils were validated using thioflavin T assays and TEM (**Figs. 1-3**). The results from mass photometry indicated that the mass of the resulting fibrils was more than 2 MDa. The fluorescence emission wavelengths of ligands in the presence of fibrils were measured by using fluospectrometer and summarized in **STable. 2**.

**Fig. 1.**
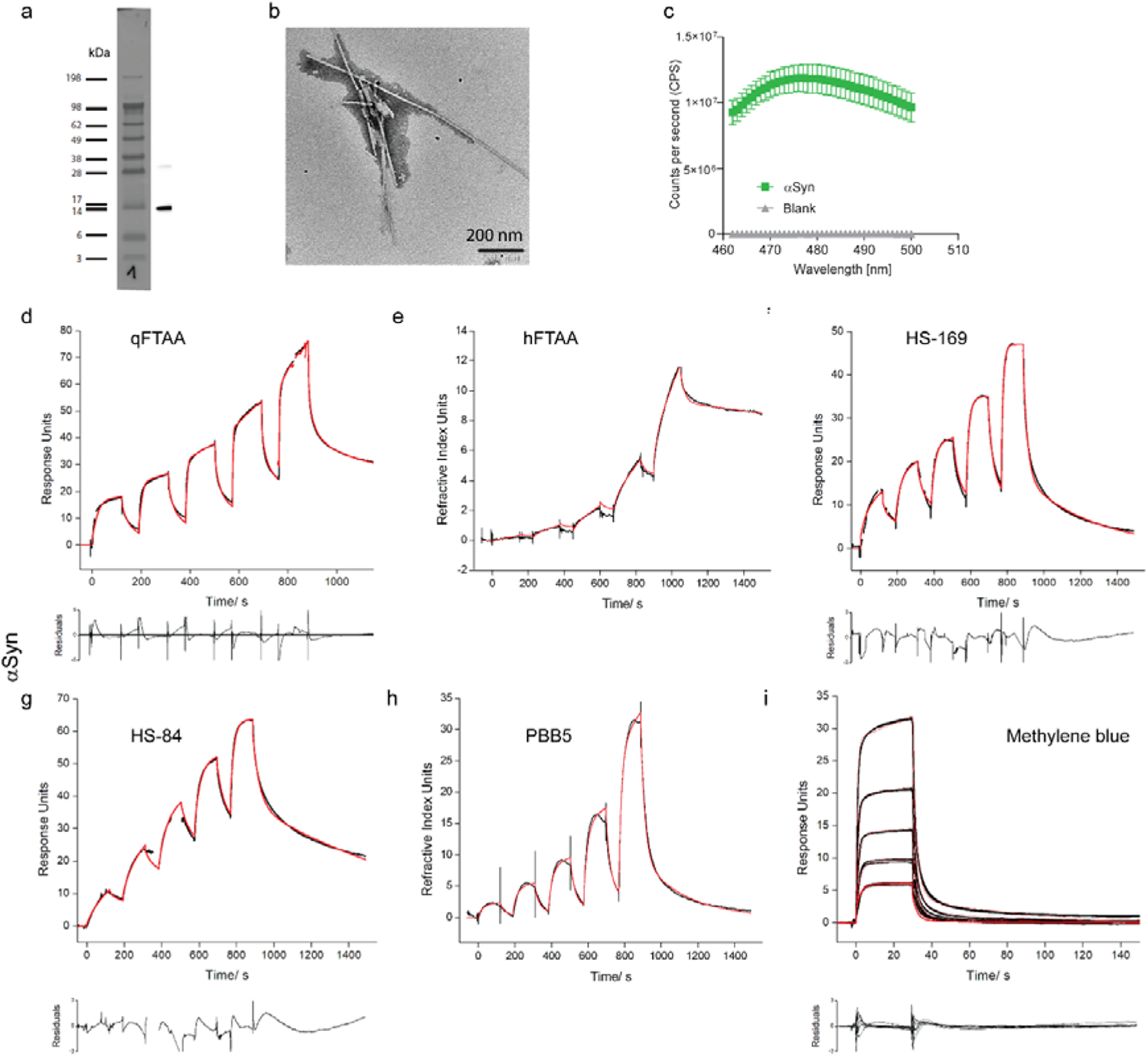
Binding data on recombinant alpha-synuclein fibrils using different surface plasmon resonance and in silico modelling of multiple binding sites. (**a**) Western blot characterization of αSyn using Anti-α-Synuclein antibody, Mouse monoclonal, clone Syn211; (**b**) TEM of αSyn fibril; (**c**) thioflavin T assay on αSyn (green) and blank (dd. water, gray) using spectrofluorometric measurements. (**d-g**) Sensorgrams of αSyn fibrils covalently immobilized on sensor chips with a 3D-hydrogel chip surface. Analytes including q-FTAA (d), h-FTAA (e), HS-169 (f), HS-84 (g), PBB5 (h), and methylene blue (i) were investigated. Black line (experimental data). Red line (fitting curve). Details on chip, chip surface, Kd evaluation in Table 1.α

**Fig. 2.**
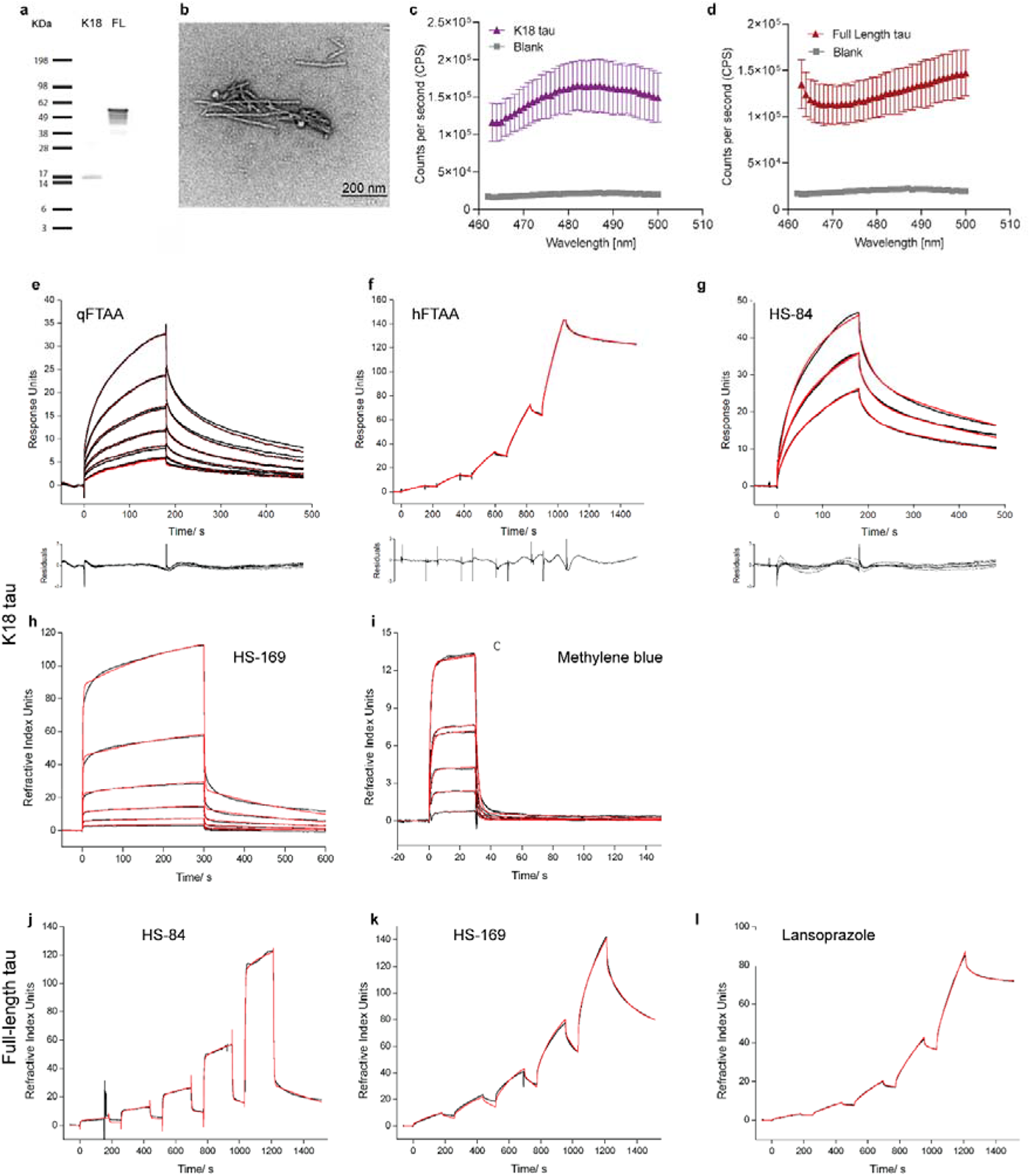
SPR binding data on recombinant K18 tau and full-length tau fibrils using surface plasmon resonance. (**a**) Western blot characterization of K18 tau and full-length tau using Anti-Tau (RD4) Antibody, clone 1E1/A6; (**b**) TEM of K18 tau and full-length tau fibrils; (**c, d**) Thioflavin T assay of K18 tau fibril (red) and full-length tau fibril (purple) and blank (dd. Water, gray) using spectrofluorometric measurements; (**e-i**) Sensorgrams of K18 tau fibrils covalently immobilized on a sensor chip with a 3D-hydrogel chip surface. Analytes including q-FTAA (e), h-FTAA (f), HS-169 (g), HS-84 (h), and methylene blue (i) were investigated. (**j-l**) Sensorgrams of full-length tau fibrils covalently immobilized on a sensor chip with a 3D-hydrogel chip surface. Analytes including HS-84 (j), HS-169 (k), and lansoprazole (l) were investigated. Black line (experimental data). Red line (fitting curve). Details on chip, chip surface, Kd evaluation in Table 1.

**Fig. 3.**
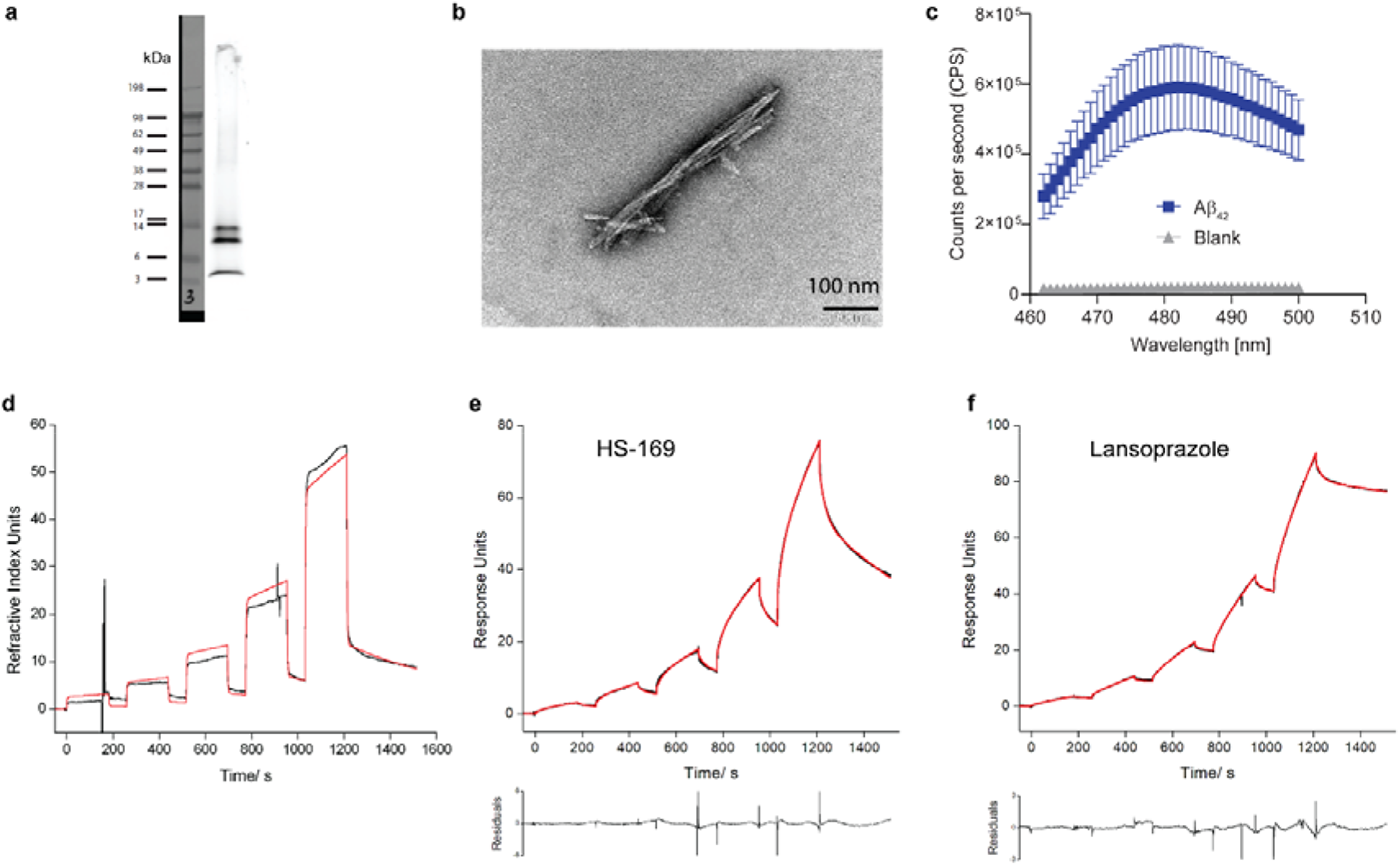
SPR binding data on recombinant Aβ42 fibrils using surface plasmon resonance. (**a**) Western blot characterization of Aβ42 using monoclonal Anti-β-amyloid antibody, clone BAM-10; (**b**) TEM of Aβ42 fibril; (**c**) Thioflavin T assay of Aβ42 fibril (blue) and blank (dd. Water, gray) using spectrofluorometric measurements; (**d-f**) Sensorgrams of Aβ42 fibrils covalently immobilized on a sensor chip with a 3D-hydrogel chip surface ZC150D. Analytes including HS-169, HS-84 and lansoprazole were investigated. Black line (experimental data). Red line (fitting curve). Details on chip, chip surface, Kd evaluation in Table 1.

### 3.2 Optimization of the SPR protocol for small molecule ligand binding to Aβ, K18 tau, full-length tau and αSyn fibrils

First, we established and optimized the experimental protocol for SPR measurement of small molecule ligands binding to Aβ, K18 tau, full-length tau and αSyn fibrils. A general problem of SPR measurements of small molecule ligands binding to immobilized fibrils is the large molar mass ratio (small molecule compound: <1000 Da, fibrils >1 MDa), which requires high surface densities to receive sufficient signal intensity. The situation is partly improved by the fact that multiple binding sites at the protein complexes lead to higher stochiometric binding ratios. Different chip surfaces were tested to achieve a high immobilization density and low nonspecific ligand adsorption. The surfaces tested included carboxymethylated dextran surface CM5, carboxymethyl dextran hydrogel surface CMD200M, linear polycarboxylate hydrogel surface HC30M and HC1500M, linear polycarboxylate hydrogel (reduced charge) surface HLC30M, and zwitterionic hydrogel surface ZC150D (**Table 2 Figs. 1-3**).

**Table 2.**
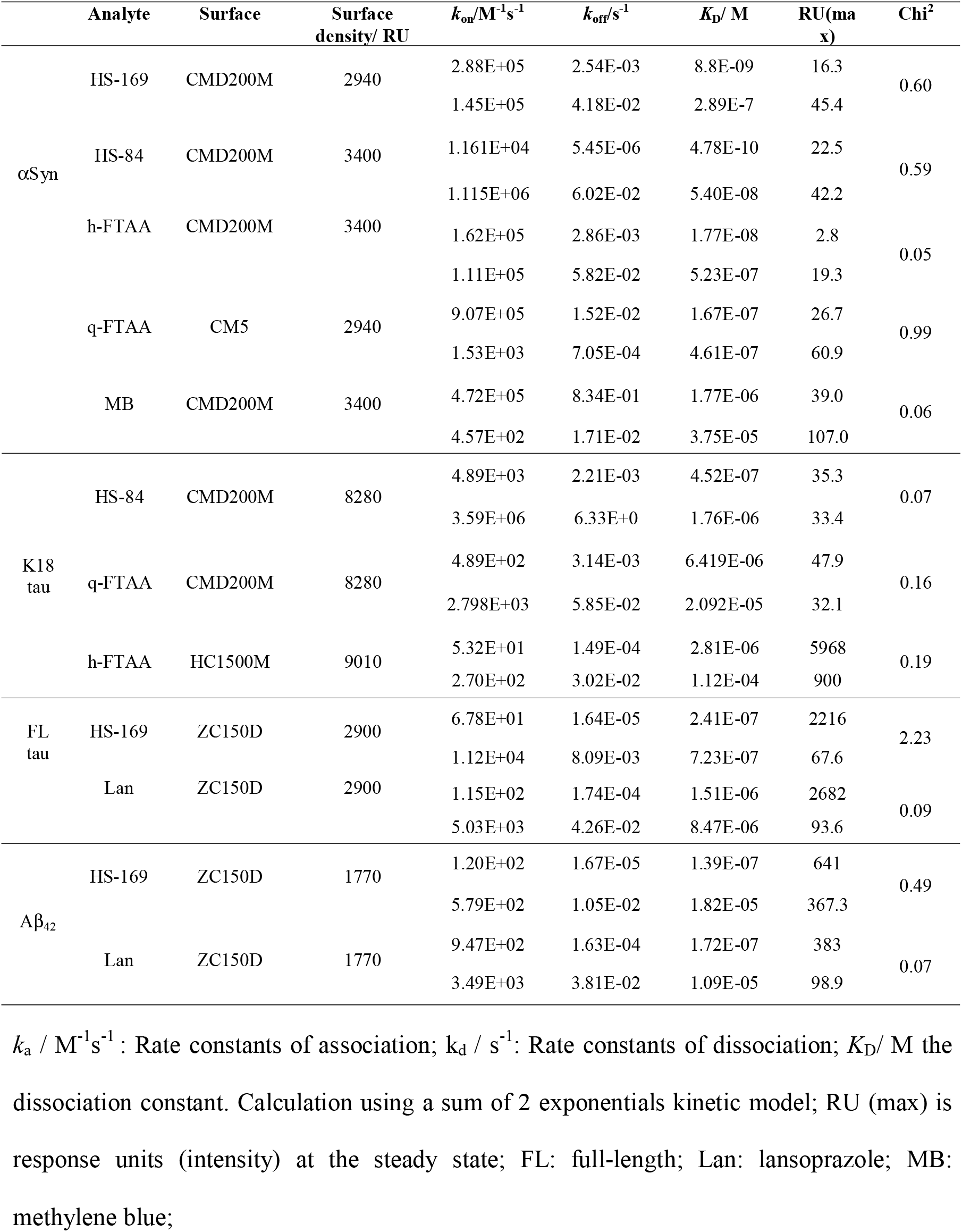
Results of surface plasmon resonance measurements of 〈Syn fibrils and A®_42_ fibrils, K18 and full-length tau fibrils.

Measurements were further complicated by the concentration of analyte solutions that usually range between approximately 1/10 of *K*_D_ and 10× *K*_D_. In the case of low-affinity binders, high nM or µM ligand concentrations often lead to strong adsorption of positively charged and hydrophobic compounds to the negatively charged chip surface, which compromises measurements. Unspecific adsorption can amount to several hundred response units (RU), thus masking binding events or making reliable quantitation impossible. As a rule of thumb, the unspecific contribution to the signal should not exceed approximately 10% of the overall signal. This can in some cases be achieved by working at very low ligand concentrations (below *K*_D_), which results in a decrease in unspecific adsorption, and the use of a surface at which adsorption is low. For this reason, in a first test, ligands were injected for 30-60 s at a concentration of approx. 1 µM to the blank chip surface to determine the adsorption behaviour.

### 3.3 Multiple binding sites of small molecule ligands to Aβ42, K18 tau, full-length tau and αSyn fibrils

The measured sensorgrams reveal complex kinetics that cannot be described by a simple 1+1 kinetic model for small molecule ligands binding to Aβ, K18 tau, full-length tau and αSyn fibrils (**Figs. 1-3**). Models of higher order, including a sum of 2 and sum of 3 exponential models, were better suited, leading in most cases to good fits. Rate constants for association and dissociation, equilibrium dissociation constants, and R_max_ values are summarized in **Table 2**. For αSyn fibrils, two binding sites (one high-affinity and one low-affinity) were observed for HS-169, HS-84, h-FTAA, q-FTAA, and methylene blue, of which HS-84 and HS-169 showed the most promising binding profiles (0.48 nM, 54 nM and 8.8 nM, 289 nM, respectively). For Aβ_42_ fibrils, two binding sites were observed for HS-169 (139 nM, 18.2 μM) and lansoprazole (172 nM, 10.9 μM) using the Sum2Exp 2 sites fitting. Although using Sum3Exp 3 sites fitting, HS-169 showed 4.1 nM, 912 nM and 9.13 μM binding site (**STable 3**). For K18 tau fibrils, two binding sites were observed for HS-84 (452 nM, 1760 nM), q-FTAA (6.41 μM, 20.9 μM) and h-FTAA (2.81 μM, 112 μM). For full-length tau fibrils, two binding sites were observed for HS-169 (241 nM, 723 nM) and lansoprazole (1.51 μM, 8.47 μM). We observed different association and/or dissociation constants and affinities of small molecules towards immobilized Aβ_42_, K18, full-length tau and αSyn fibrils by SPR. The *k_on_* and *k_off_* values fit with the observation of slow wash-out features of HS-169 and HS-84 reported *in vivo* (Calvo-Rodriguez et al., 2019b; Ni et al., 2022; Ni et al., 2020).

### 3.4 In silico modelling demonstrated multiple binding sites of LCOs on αSyn fibrils

Here, we used the 6H6B structure from the recombinant αSyn fibril, which is assembled in paired helical fibril (PHF) form. From the molecular docking studies of HS-169, HS-84, p-FTAA, and q-FTAA, we identified four binding sites on the αSyn fibril (6H6B, **Figs. 4a, b**), denoted as, sites 1-4. The salt bridge between E57 and H50 and the van der Waals interaction between the shallow hydrophobic residues (G51, A53, and V55) is found to stabilize the paired fibril in 6H6B. Site 3 is a core site inside the fibril and can be easily accessed when a limited number of molecules are used in the modelling, which might not be easily accessible in real situations. We carried out a 100-ns MD simulation for the binding of each ligand on each site, resulting in 20 trajectories with a total length of 2 μs. The ligands HS-169, HS-84, p-FTAA and q-FTAA are negatively charged; therefore, the binding affinities at site 4, which is a positively charged site, are more favourable than those at site 3 (**Fig. 4c**). In line with the docking studies, the ranking of binding free energies indicates that HS-169 and HS-84 show very strong binding at site 4, followed by p-FTAA, q-FTAA and h-FTAA. These compounds are rich in hydrogen acceptors/donors and form stable hydrogen bonds with a cluster of positively charged residues K43, K45, H50, and K58 (**Fig. 4c**). The ligand RMSD to the initial conformation, usually used to estimate the stability of the binding, shows a deviation of larger than 8Å, which indicates that the ligand has moved away from the original binding site (**Figs. 4d-g**). From this point of view, the stability of the binding sites can be ranked as S4 > S3 > S2 > S1, for the binding the five investigated ligands. Additional in silico modelling was performed using the 2N0A αSyn structure. Four binding sites for DCVJ have been demonstrated on the 2N0A αSyn structure (Kuang et al., 2019). Here, we found a preference of these ligands for the core site 3 using the 2N0A α yn structure (**SFigs. 3**).

**Fig. 4.**
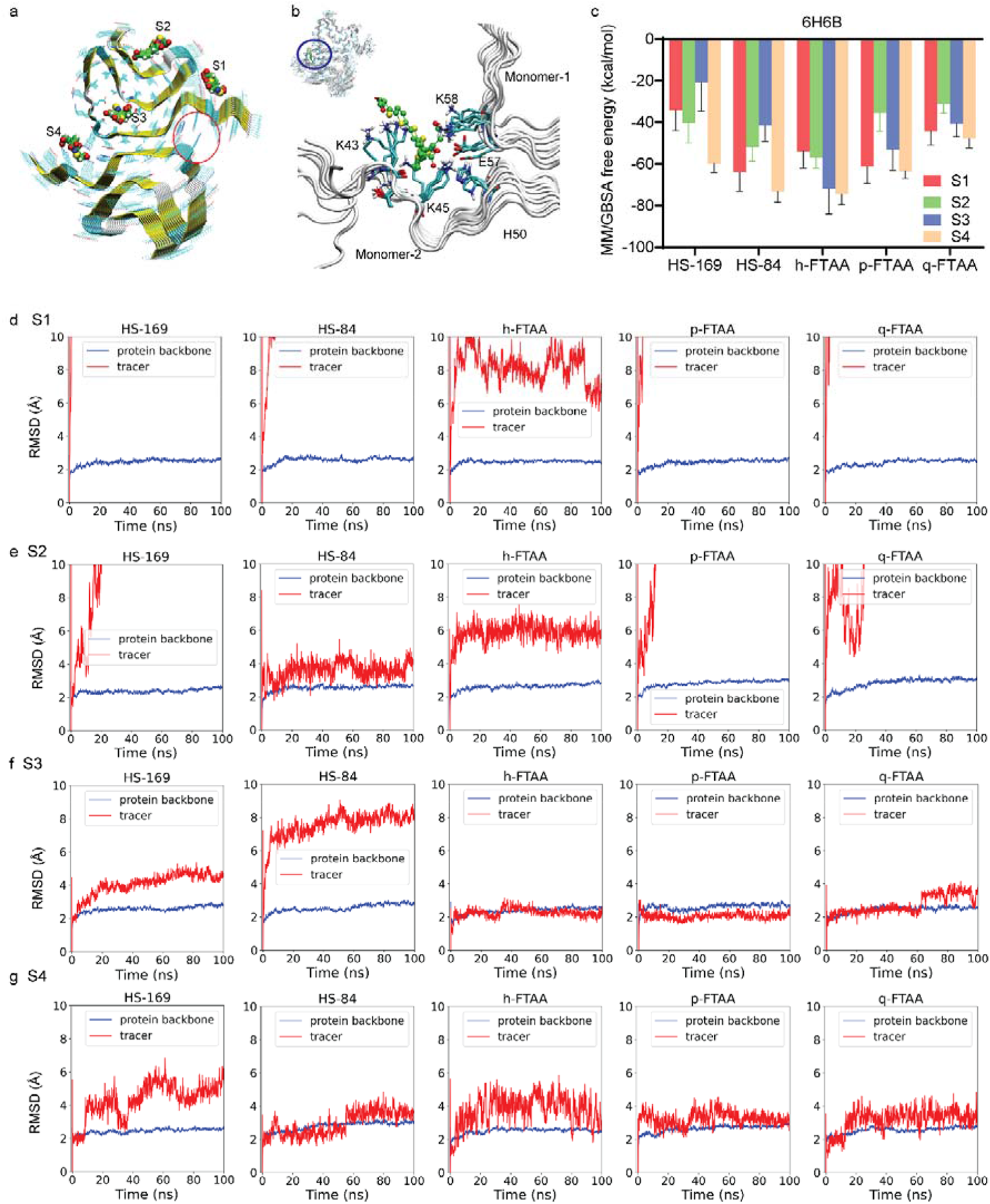
In silico modelling of the binding sites of HS-169, HS-84, h-FTAA, p-FTAA and q-FTAA on the 6H6B alpha-synuclein structure. (**a**) Four binding sites (S1-S4) on alpha-synuclein fibrils; the red circle indicates the location of another site 4; (**b**) Zoomed-in view of h-FTAA binding to site 4. (**c**) MM/GBSA calculation of free energy indicating that site 4 is preferred by HS-169, HS-84, h-FTAA, p-FTAA and q-FTAA. (**d-g**) RMSD analysis of the ligands binding to 4 binding sites (S1-S4).

### 3.5 Validation using immunofluorescence staining of postmortem human brain tissue and mouse models

We investigated the ligand detection of αSyn inclusions using the phospho-αSyn antibody pS129. We showed that ligands (h-FTAA, q-FTAA, and HS-169) were bound to positive pS129 αSyn inclusions in medulla oblongata and locus coeruleus tissue slices from three patients with PD (**Figs. 5a-e**). No positive signal in the non-demented control was observed (**Fig. 5f**). Colocalization of different ligands to both the Lewy neurites and Lewy bodies were observed. Next we evaluated the detection of HS-169 and HS-84 in αSyn-PFF-injected mice at 12 weeks after injection into the pedunculopontine nucleus (PPN). Positive pS129 signal was detected in the ChAT-positive cholinergic PPN neurons of αSyn-PFF-injected mice (**Fig. 5g**). Colocalization of the ligands (HS-169 and HS-84) with the pS129 signal was observed in the periaqueductal gray, nucleus accumbens, and central amygdala of αSyn-PFF-injected mice (**Figs. 5g-j**). The somatic pS129-positive αSyn aggregates appear to be better stained by ligands compared to the neuritic pS129-positive αSyn pathology (**Fig. 5i**).

**Fig 5.**
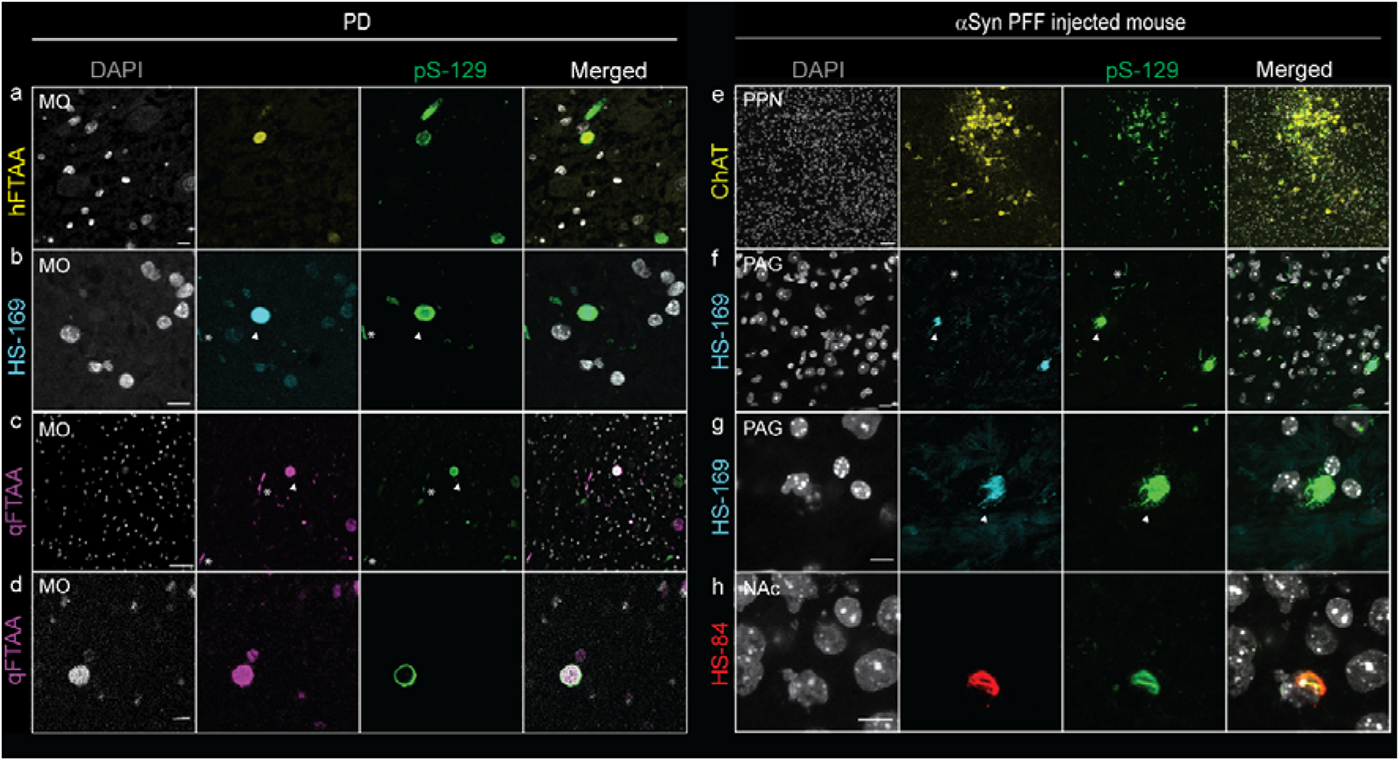
Confocal imaging of alpha-synuclein inclusions in brain tissue sections from PD patients and αSyn PFF-injected mice. (**a-d**) Immunofluorescence staining in the medulla oblongata of PD patients. αSyn-positive Lewy neurites (*) and αSyn-positive Lewy body (arrowhead). Colocalization of Alexa488-anti-aSYN phosphor S129 antibody pS129 (green), with h-FTAA (yellow), HS-169 (cyan), and q-FTAA (magenta); Nuclei were counterstained using DAPI (white). (**e-h**) Immunofluorescence staining in αSyn PFF-injected mouse brain. pS129-positive inclusions in ChAT-positive neurons (yellow) in the PPN of αSyn PFF-injected mice. (**f-g**) Colocalization of pS129-positive inclusions (green) with the ligand HS-169 (cyan) in the periaqueductal gray matter (PAG, h is Zoom-in-view of g). **(h**) Colocalization of pS129-positive inclusions (green) with HS-84 (red) in the nucleus accumbens (NAc); nuclei were counterstained using DAPI (white). Scale bar = 50 μm (c, e), 10 μm (a, b, d, f, g, h).

To validate the detection of Aβ or tau pathology, we studied the binding of the aforementioned ligands to transgenic mouse brains with Aβ or tau pathology. We chose transgenic mouse models to study Aβ or tau pathology since the brain tissue from AD patients commonly shows coexistence of these two pathologies. Tau mainly accumulates in the cortex and hippocampus of pR5 mice. Immunofluorescence staining performed on coronal brain tissue sections from pR5 and nontransgenic mice co-stained with anti-phospho-tau AT-8 antibody in the cortex and hippocampus confirmed the detection of PBB5 to tau inclusions (**Figs. 6a-c**). AT-100 antibody (mature phospho-tau) staining with PBB5 was performed in brain tissue slices from pR5 mice and further validated the detection (**Figs. 6a, b**).

**Fig. 6.**
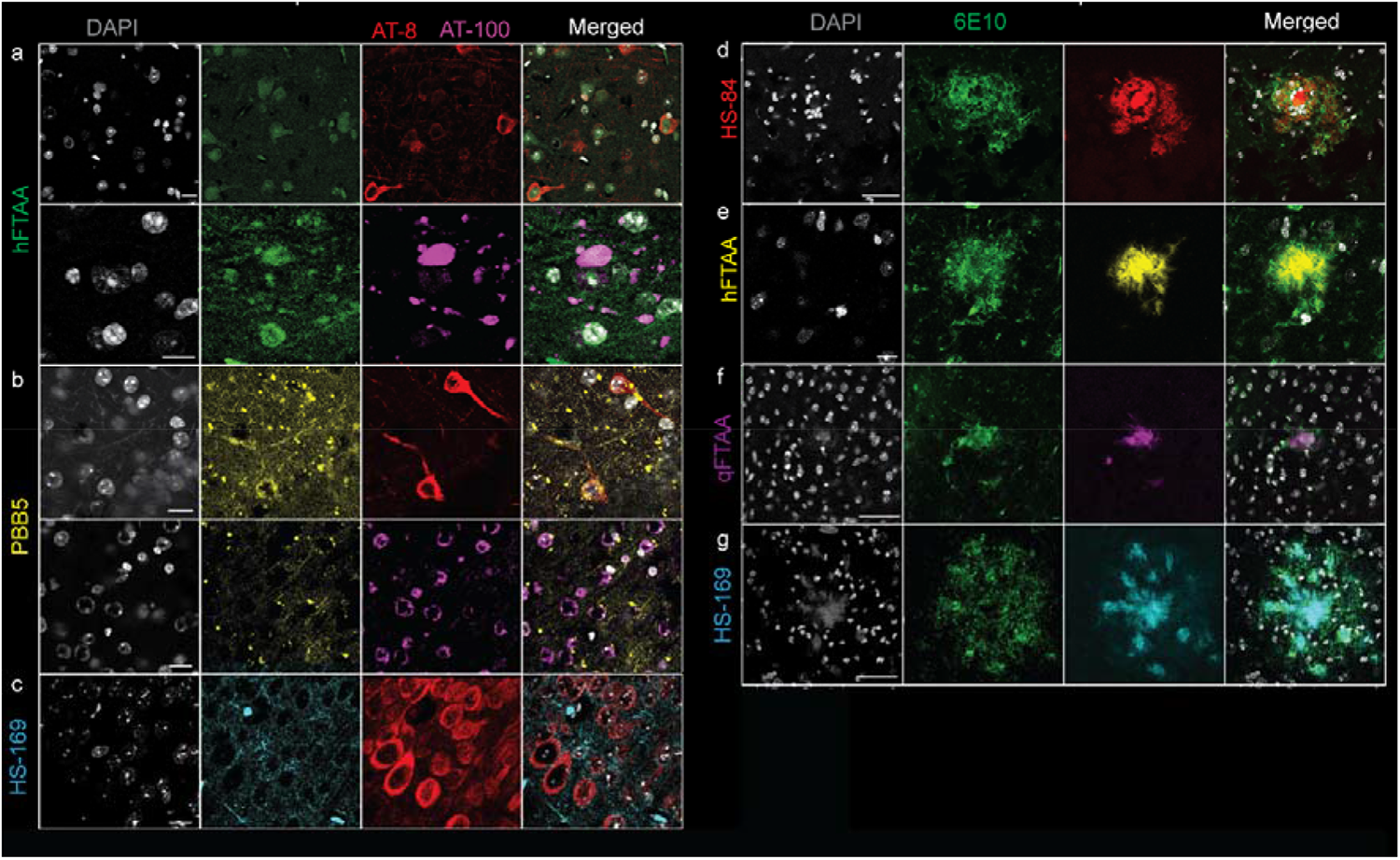
Colocalization of imaging ligands with tau and amyloid-beta. (**a-c**) Immunofluorescence staining in the hippocampus of pR5 tau mice. Alexa488-AT8 (red), Alexa488-AT100 (magenta), h-FTAA (green), PBB5 (yellow), HS-169 (cyan); nuclei were counterstained using DAPI (white). Scale bar = 10 μm. (**d-g**) Amyloid-beta deposits in the cortex of arcAβ mouse brain tissue sections. Anti-Aβ1-16 antibody Alexa488-6E10 (green), HS-84 (red), h-FTAA (yellow), q-FTAA (magenta), HS-169 (cyan). Nuclei were counterstained using DAPI (white). Scale bar = 50 μm (d, f, g), 10 μm (a, b, c, e).

We assessed ligand detection of Aβ deposits in brain tissue slices from arcAβ mice using 6E10. arcAβ mice develop Aβ pathology affecting both the brain parenchyma and vasculature from 6 months of age (Kecheliev et al., 2022; Knobloch et al., 2007). Immunofluorescence staining performed on coronal brain tissue sections from arcAβ and nontransgenic mice with ligands (h-FTAA, q-FTAA, and HS-169) co-stained with 6E10 (anti-Aβ_1−16_) antibody further confirmed the detection of parenchymal Aβ deposits in the cortical regions (**Figs. 6d-g**).

## 4 Discussion

In the present work, we optimized the SPR approach to characterize the multiple binding sites of imaging ligands, and small molecule to Aβ_42_, K18 tau, full-length tau and αSyn fibrils. SPR measurements indicate that, for all fibrils, the binding kinetics of the imaging ligands and small molecules are complex since the sensorgrams recorded cannot be evaluated by fitting a simple 1+1 kinetic model. Kinetic models of higher order were better suited to fit the data, including a sum of 2 and sum of 3 exponential models. On the assumption that ligand binding to one of the 4 sites calculated for αSyn (**Fig. 4**) takes place independently, each interaction is characterized by a specific association and dissociation rate constant, resp., and relative weight. In this case, the order of the fit model should correspond to the number of binding sites. Indeed, models of higher order, in particular the sum of 3 exponentials model, considerably improved fit results as judged by chi^2^ and the distribution of residuals (**Table 2**).

The fit results confirm the existence of multiple binding sites in agreement with modelling calculations. However, it needs to be noted that fitting data with models of higher order introduces considerable uncertainty due to the large number of initial parameters (*k*_a_, *k*_d_, R_max_ for each additional interaction: A sum of 2 (3) exponential models has at least 6 (9) free fit parameters plus (optional) a parameter for bulk response (RI) and correction (t_c_). This can cause the fit routine to become stuck in local minima. This possibility can be shown by repeating a fit with a single parameter being changed by a larger value (two orders of magnitude), a procedure that often does not reproduce the initial fit. Extensive tests with the sum of 3 exponentials model revealed that fit data show low resistance to changes in the initial parameters, and the results should be interpreted with caution. The number of exponentials needed to fit a sensorgram does not necessarily correlate with the number of binding sites identified in modelling calculations. It represents its lower limit since some interactions may not be resolved due to the similarity of rate constants and a general limit of data quality of SPR measurements. For this reason, we limited the evaluation of sensorgrams to the sum2exp model, although for the sum3exp model, chi2 was often found to be much lower, and residuals showed a more random distribution without a recognizable pattern. In the case of sensorgrams of multiple and complex interactions, calculated rate constants should be interpreted with caution.

The binding of different ligands on the various sites of these fibrils with abnormal ligand binding cavities has also been evaluated by free energy calculations to rank the affinities. In silico modelling suggested 4 binding sites on αSyn with a preference for certain binding pockets of different molecules (**Fig. 4**). The stabilities of the binding sites identified by docking were further evaluated by MD simulations and binding free energy calculations. These studies provide atomic details as well as identify key amino residues for the interaction between ligands and fibrils. One here often witnesses an induced-fit mechanism of ligand binding, e.g., enlargement of a narrow cleft in the protofibril when a large and planar ligand binds. Planarity of the ligand tends to increase its van der Waals or π-π interactions with the β-sheets of the protein and, due to a loss of molecular flexibility, an increase in fluorescence quantum yield (LIT). The core site, which is buried inside the protofibril, can often exhibit high affinities to small molecules due to enhanced van der Waals interactions. However, here we observed a surface site with strong binding affinities to the negative charged ligands due to the ionic interactions, indicating that the way the fibril twists is crucial for the existence of a site (S4) surrounded by several base residues (K43, K45, H50, and K58). In addition to the binding free energies, the residence time of the ligands at the binding sites of fibrils is also relevant to consider to rank the potency of the ligands, which can be estimated by the dissociation rate constant, *k*_d_, measured with SPR. If a ligand can preferentially bind to the core site(s) inside the protein fibrils, as in the presently studied case, there is usually stronger binding affinity and longer wash-out time. Several recent studies have suggested 6 binding sites on Aβ fibrils (Arul Murugan et al., 2018; Balamurugan et al., 2017; Kawai et al., 2018; Kuang et al., 2015; Murugan et al., 2016; Murugan et al., 2014) and at least 4 binding sites on tau fibrils (Kuang et al., 2020; Murugan et al., 2019; Murugan et al., 2018; Murugan et al., 2021). Kuang et al showed several surface and 3 core binding sites of PI-2620 (Kuang et al., 2020) on tau fibrils and one core and 3 surface sites on αSyn fibrils (Kuang et al., 2019).

Our affinity data of LCOs were in line with available binding data towards Aβ_42_ from previous studies. Johansson et al. immobilized azide-functionalized p-FTAA and showed a K_d_ of ∼10 nM towards Aβ_42_ using SPR as well as ratiometric comparison of the excitation spectra for free vs. bound dye (Johansson et al., 2015). Using competition studies with [^3^H]X-34, Bäck et al. showed that h-FTAA and q-FTAA bound to recombinant Aβ_42_ fibrils with EC50 ∼250 nM and 330–630 nM (Bäck et al., 2016). In addition, Herrmann et al. showed ratiometric comparison of the excitation spectra for free versus bound dye to determine K_d_ values for some LCOs bound to recombinant PrP fibrils, and the K_d_ values were in the low nM range (Herrmann Uli et al., 2015).

Autoradiography and in situ binding assays on brain tissues have also been used for evaluating the binding affinity, but in general, only one binding site can be obtained (Han et al., 2011). For fluorescent probes, radiometric analysis has been used to estimate the binding sites with less accuracy. Using radioligand binding assays, multiple ligand binding sites on Aβ fibrils (LeVine, 2005; Ni et al., 2017; Ni et al., 2013a; Ni et al., 2021a) and tau fibrils (Malarte et al., 2020; Ni et al., 2018; Yap et al., 2021), e.g., using THK-5351 (Lemoine et al., 2017; Lemoine et al., 2020; Stepanov et al., 2017), MK-6240 (Lemoine et al., 2021; Malarte et al., 2022; Malarte et al., 2020), and PBB3 (Ono et al., 2017), have been reported. Different tau ligands show distinct binding towards tau in AD and different primary tauopathy (Shi et al., 2021a; Yap et al., 2021). A recent study observed a drug candidate binding to the core site of αSyn from a cryo-EM density map (Antonschmidt et al., 2022).

In addition to the affinity information, rate constants for different compounds vary by orders of magnitude for the association from (10E^1^-10E^9^ M^-1^ s^-1^) and 0.1 s^-1^ to 1E^-8^ s^-1^ for the dissociation step (kinetic data in **Table 2**). The calculated equilibrium constants *K*_D_ are mainly in the nM to µM range, in agreement with the fact that these dyes have been successfully used in fluorescence imaging applications (Calvo-Rodriguez et al., 2019a; Ni et al., 2022; Ni et al., 2020). Some targets have components in the pM range when dissociation is very slow, including αSyn, K18 tau fibrils with h-FTAA and Aβ_42_ fibril with lansoprazol. For the αSyn fibril, it is obvious to assign the components found in the analysis of sensorgrams to the binding sites calculated with molecular mechanics. In such a case, fast binding is expected to take place at sites that are easily accessible, e.g., site 1 and site 4 in **Fig** In contrast, slow association and dissociation components are attributed to core site 3, which is more difficult to access. For K18 tau and Aβ_42_ fibrils, no MM calculations are available. The similarity of sensorgrams between αSyn suggests that multiple binding sites should also exist in these fibrils. In light of the difficulty in comparing rate constants calculated with higher-order kinetic models, a classification of targets based on binding properties of the main component may be useful. Basically, there are two types of sensorgrams depending on whether the interaction is governed by a fast (Type 1) or slow (Type 2) association and dissociation (**Table 2**). In the case of slow dissociation and assuming that the interaction *in vivo* follows the same kinetics, the dye is most likely suitable for imaging applications. As a consequence, screening by SPR can be useful to assess whether a target is suitable for imaging applications.

We further demonstrated the colocalization of different fluorescence-emitting probes on brain tissue slices from PD patients, αSyn PFF-injected mice, and transgenic mice with Aβ plaques or tau inclusions, in line with previous observations (Ni et al., 2020; Ni et al., 2021b; Ren et al., 2022). The LCOs tested here appear to detect the αSyn inclusions in the αSyn PFF-injected mouse brain and can be useful in the *in vivo* imaging studies.

There are several limitations of our study. First, we did not use brain tissue-derived fibrils from patients with AD, primary tauopathy, PD, and MSA to investigate the ligand binding profiles. The binding sites on recombinant fibrils might differ from those on fibrils derived from patients (Lövestam et al., 2021). Further studies using fibril pulldown from human brain tissue for potential strain-dependent affinity will provide important insight because LCOs have been shown to discriminate αSyn strains in PD and MSA (Klingstedt et al., 2019; Shahnawaz et al., 2020). Second, we did not measure the binding to oligomeric forms of Aβ, tau, and αSyn by SPR due to the uncertainty of oligomer status on chips as the fibril forms rapidly. Regarding fitting of SPR data, a sum of 2 exponential models to adjust the model to the number of binding sites found in calculations resulted in data of little confidence, which is caused by the large number of free fitting parameters. Therefore, it was reasonable to expect that fitting with a sum of 4 exponential kinetic models would not lead to useful fit data.

## 5. Conclusion

In this work, we developed a binding property characterization platform for small molecule binding to Aβ_42_, 4R tau, full-length tau and αSyn fibrils. Such a system will greatly improve the efficiency of imaging ligands and drugs targeting these protein fibrils, which are critical for the diagnosis and treatment of neurodegenerative diseases such as AD, PD, and tauopathies. In addition, such a platform will also be potentially useful for studying competition of binding sties and off-target binding by displacement assays.

## Supporting information

SFig, STable

## 6 Declaration

### Authors’ contributions

The study was designed by RN. JS conducted SPR measurements and kinetic analysis. JL and HA performed and analysed the in silico modelling. JG performed the fibril production and fluorescence binding studies. BC performed staining and confocal microscopy. MH and FG provided the αSyn PFF-injected mouse brains. KPN provided the LCOs. JS, JL, and RN wrote the first draft. All authors contributed to the revision of the manuscript. All authors read and approved the final manuscript.

## Acknowledgements

The authors acknowledge the Center for Microscopy and Image Analysis (ZMB), University of Zurich; Dr. Saroj Kumar Rout, Mr Patrick Vagenknecht, ETH Zurich, Prof. Jan Klohs, Institute for Biomedical Engineering, ETH Zurich/University of Zurich, and Dr. Uwe Roder at Cytiva Europe GmbH.

## Supplemental Material

**SFig 1.**
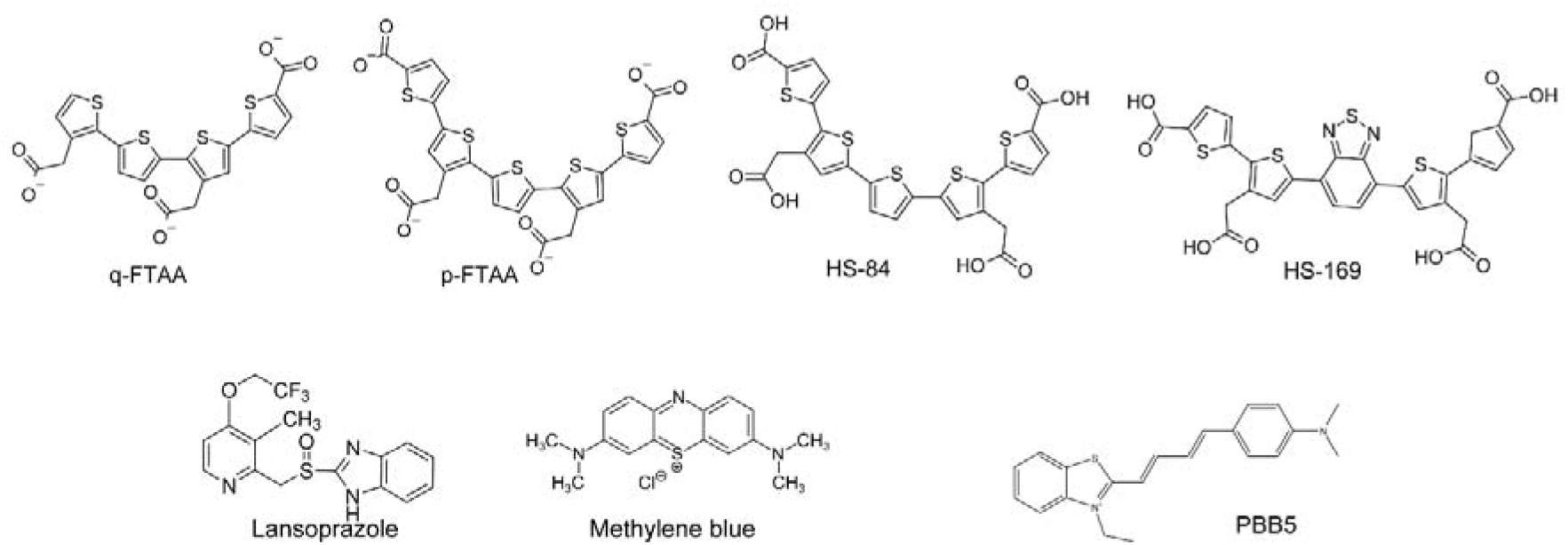
Chemical structures of the ligands and compounds.

**SFig 2.**
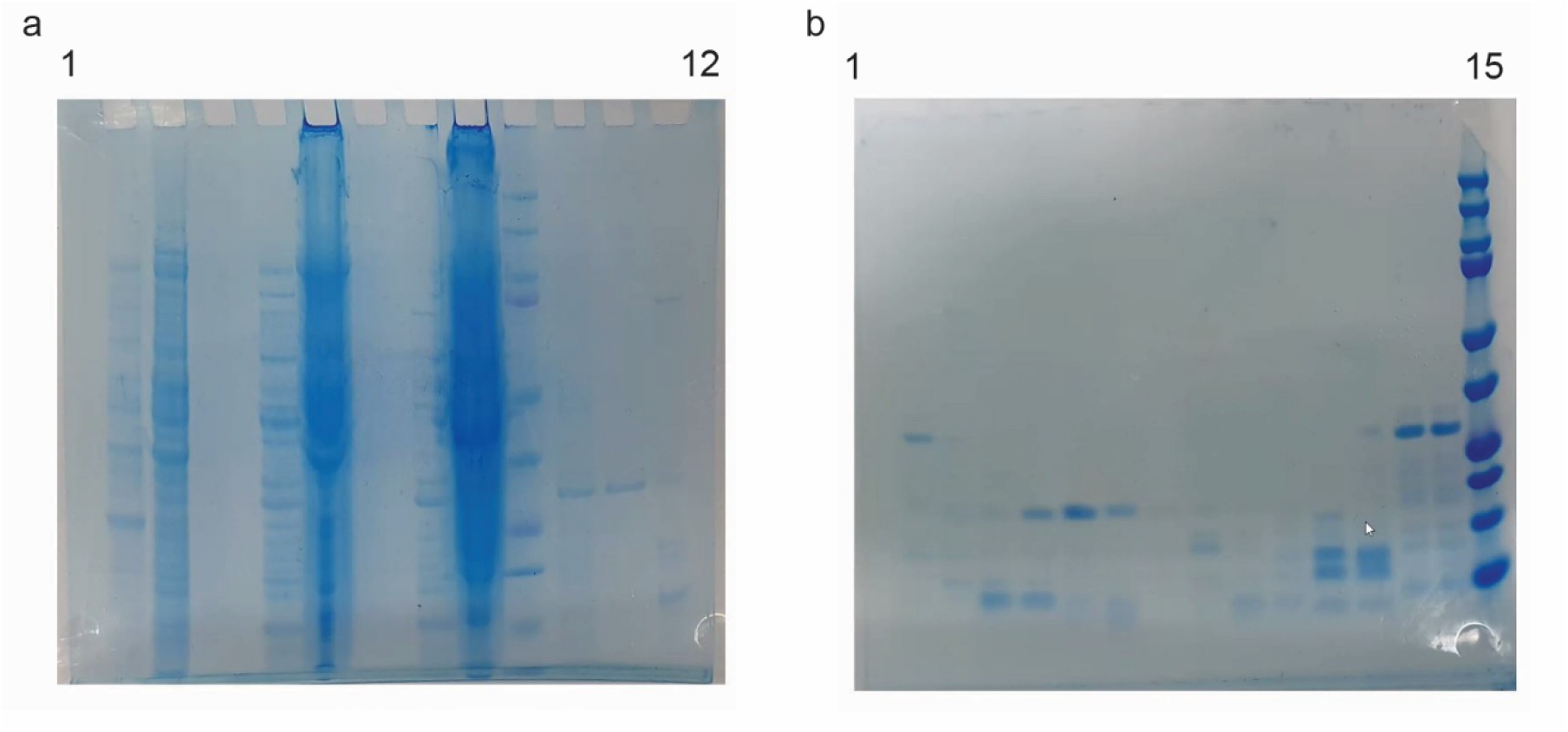
Confirmation of αSYN and K18 tau. (**a**) αSYN purification (bands 1-12). After each step, an aliquot was taken, and ultimately, the identity was verified by sodium dodecyl sulfate–polyacrylamide gel electrophoresis (SDS–PAGE). The identity of αSYN was confirmed by a visible band at approximately 14 kDa (in band 12). From bands 1-12 (left to right): 1) *E. coli* uninduced (before adding IPTG), 2) *E. coli* induced, 3) empty, 4) osmotic shock (OS) supernatant (Sup.), 5) OS pellet, 6) empty, 7) heat shock (HS) supernatant, 8) HS pellet, 9) Bio-Rad Precision Plus Protein Dual Color Standards, 10) ammonium sulfate (AS) precipitation 35% supernatant, 11) AS 55% supernatant, 12) AS 55% pellet. (**b**) tau purification (bands 1-15). After centrifugation of the cell debris, the supernatant was recovered, and column chromatography with a phosphocellulose column was performed. Twelve elution fractions of the chromatography were obtained and verified by SDS–PAGE (bands 1-12, left to right). The three fractions with the highest Tau concentrations (fractions 4, 5, and 6) were selected for subsequent dialysis and lyophilization. Bands 13-15 were the solution before chromatography, the flow-through, and the reference band (Bio-Rad Precision Plus Protein Dual Color Standards).

**SFig 3.**
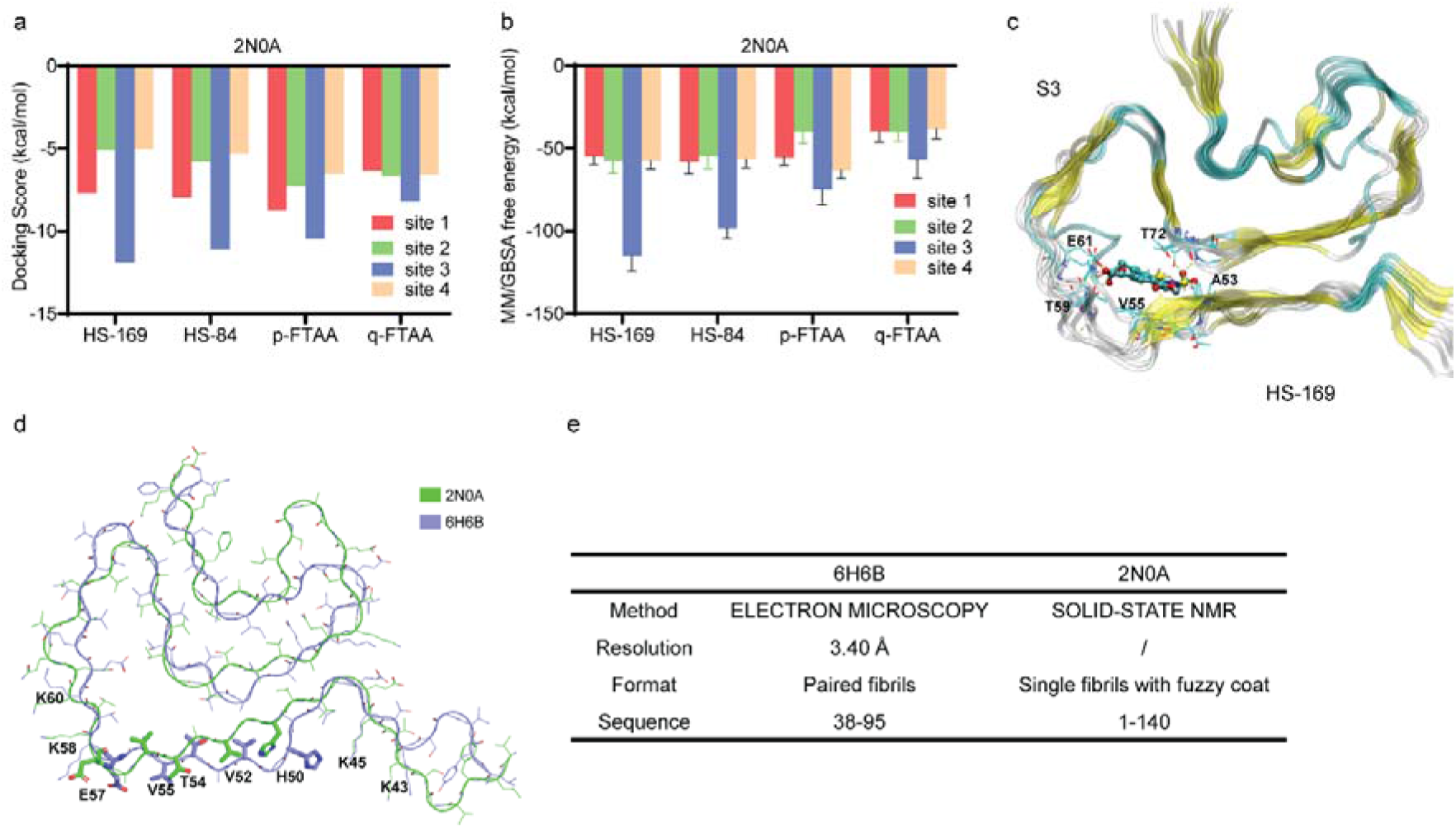
In silico modelling of the binding sites of HS-169, HS-84, h-FTAA, p-FTAA and q-FTAA on the 2N0A alpha-synuclein structure. **(a, b)** Docking and MM/GBSA calculation of free energy indicating that site 3 is preferred by HS-169, HS-84, h-FTAA, p-FTAA and q-FTAA on the 2N0A alpha-synuclein structure. **(c)** Zoomed-in view of HS-169 binding to site 3. **(d, e)** Difference between the 2N0A and 6H6B structures.

