## Supplementary material for "Efficient characterization of multiple binding sites of small molecule imaging ligands on amyloid-beta, 4-repeat/full-length tau and alpha-synuclein": SFig, STable

**Supplemental Table 1** **List of compounds in the fluorescence assay, SPR assay and primary antibodies in staining**

| **Ligands/Compounds** | **Source** |  | | | **Conc [μM] in staining** |
| --- | --- | --- | --- | --- | --- |
| PBB5 | RadiantDye, |  | | | 1.6 |
| Methylene Blue | Sigma‒Aldrich | | |  |  |
| HS-169 | KPRN | | |  | 5 |
| HS-84 |  |  |  |  | 5 |
| h-FTAA |  |  |  |  | 5 |
| q-FTAA |  |  |  |  | 5 |
| Lansoprazole | Sigma‒Aldrich | |  | |  |
| **Antibody/Compounds** | **Source** | | | **Cat. No.** | **Dilution** |
| DAPI | Sigma‒Aldrich | | | D9542-10MG | 1:1000 |
| Goat-anti-Rabbit Alexa488 | Invitrogen | | | A11034 | 1:200 |
| 6E10, Anti-β-amyloid, 1-16 antibody | Biolegend | | | 803001 | 1:1000 |
| AT-8, anti-Phospho-tau Ser202, Thr205 | Invitrogen | | | MN1020 | 1:1000 |
| AT-100, anti-Phospho-tau Thr212, Ser214 | Invitrogen | | | MN1060 | 1:1000 |
| Anti-a-synuclein pS129 (phospho-Ser129) | Abcam | | | ab51253 | 1:1000 |
| Donkey anti-Rabbit Cy5 | Jackson ImmunoResearch | | | 711-175-152 | 1:200 |
| VECTASHIELD® Antifade | Vector Laboratories | | | H-1000-10 |  |
| Anti-Choline Acetyltransferase Antibody | Sigma‒Aldrich | | | AB144P-1ML5 | 1:100 |
| Normal goat Serum (NGS) | Chemie Brunschwig | | | JAC005-000-121 |  |
| Normal Donkey Serum (NDS) | Interchim-Uptima | | | UP77719A K |  |
| Triton X-100 | Sigma‒Aldrich | | | X100-500ML |  |
| Anti-Tau (RD4) Antibody, clone 1E1/A6 | Sigma‒Aldrich | | | 05-804 | 1:1000 |
| Anti-α-Synuclein antibody, Mouse monoclonal, clone Syn211 | Sigma‒Aldrich | | | S5566 | 1:2000 |
| Monoclonal Anti-β-amyloid antibody, clone BAM-10 | Sigma‒Aldrich | | | A5213 | 1:1000 |
| Anti-mouse IgG antibody conjugated to horseradish peroxidase (HRP) | Jackson ImmunoResearch | | | 115-035-166 | 1:5000 |
| ECL Prime Western Blotting Detection Reagents | Cytiva | | | RPN2232 |  |
| Donkey-anti-Rat Alexa488 | Jackson ImmunoResearch | | | AB2340686 | 1:400 |
| Goat-anti-mouse Alexa488 | ThermoFischer Scientific | | | A11001 | 1:200 |

**Supplemental Table 2 Fluorescence emission wavelengths of ligands in the presence of fibrils**

| **Ligand** | **Excitation wavelength [nm]** | **Emission wavelength [nm]** | | | | |
| --- | --- | --- | --- | --- | --- | --- |
|  |  | Aβ_42_ | K18 tau | | α Syn | |
| PBB5 | 630 | 695 | | 688 | | 690 |
| HS-169 | 375, 535 | 650 | | 640 | | 643 |
| HS-84 | 430 | 504, 540 | | 504, 540 | | 504, 540 |
| h-FTAA | 480 | 544, 573 | | 544, 577 | | 544, 577 |
| q-FTAA | 430 | 473, 502, 533 | | 473, 502, 533 | | 473, 502 |
| Methylene blue | 680 | - | | - | | - |

MW, molecular weight. The second peak of h-FTAA and HS-84 differs between binding to Aβ_42_, K18 tau and αSyn fibrils. The third peak (shoulder) of q-FTAA was more apparent in the binding to Aβ_42_ fibrils compared to K18 tau fibrils and was missing in the binding to αSYN fibrils.

**Supplemental Table 3 Result of 3 sites fitting using Sum3Exp 3 sites for surface plasmon resonance data of HS-169 binding on αSyn fibrils**

| Analyte | Surface | *k*_on_/M^-1^s^-1^ | *k*_off_/s^-1^ | *K*_D_/ nM | RU (max) | Chi^2^ | tc |
| --- | --- | --- | --- | --- | --- | --- | --- |
| HS-169 | ZC150D | 1.52E+03 | 1.38E-03 | 9.12E-07 | 163.7 | 0.321 | 3.5E+05 |
|  |  | 1.92E+05 | 7.92E-04 | 4.12E-09 | 33.8 |  |  |
|  |  | 5.83E+03 | 5.32E-02 | 9.13E-06 | 158.1 |  |  |

**Supplemental Methods**

**Fibril production**

To induce fibrillization, the lyophilized proteins were dissolved in phosphate-buffered saline (PBS) buffer pH 7.4 (Gibco, U.S. containing 0.05% NaN_3_ (Sigma Aldrich, U.S.A.). The proteins were resuspended in 200 μl of PBS/NaN_3,_ and several aliquots were prepared in 1.5 ml Eppendorf tubes. For fibrilization, the protein solutions were incubated at 37 °C under agitation (500-700 RPM) in an Eppendorf thermomixer. The proteins were incubated at the following concentrations: Aβ_42_ (50 μM), αSyn (250 µM), and K18 tau (50 µM). The gel (Novex 10-20%, Tricine, Thermo Fisher Scientific, U.S.A.) was loaded with 10 μg of K18 tau and 10 μg of αSyn. For Aβ_42_, the quantity was 3.7 μg because of a higher dilution. The samples were mixed with Laemmli sample buffer and boiled at 95 °C for 5 minutes. Sodium dodecyl sulfate–polyacrylamide gel electrophoresis (SDS–PAGE) was performed at 100 V for 1 h 45 min. The gel was transferred to a nitrocellulose membrane (Invitrogen iBlot Transfer Stack, nitrocellulose, Thermo Fisher Scientific) using a dry blotting system (Invitrogen iBlot 2, Thermo Fisher Scientific, U.S.A.). The membrane was washed with 1 × PBS and 0.1% Tween 20 (PBST) and blocked with 5% fat-free milk. K18 tau was detected using a monoclonal tau antibody (anti-tau 4-repeat isoform RD4, clone 1E1/A6). Aβ_42_ was detected with a monoclonal Aβ antibody (clone BAM-10), and αSyn was detected using a monoclonal αSyn antibody (clone Syn211). The primary antibodies were diluted in 5% fat-free milk. As a secondary antibody, a purified anti-mouse IgG antibody conjugated to horseradish peroxidase (HRP) was used (Jackson ImmunoResearch) diluted in 5% fat-free milk. The membranes were incubated with the primary antibodies at 4 °C overnight. The membranes were washed with PBST, subsequently incubated with secondary antibodies at room temperature for 2 h and then washed with PBST.
